# Yield of Universal Testing for DNA Mismatch Repair Protein Deficiency in Colorectal Carcinoma From an Australian Community-based Practice

**DOI:** 10.1101/270322

**Authors:** Gregory C. Miller, Mark L. Bettington, Ian S. Brown, Christophe Rosty

## Abstract

Lynch syndrome is the most common cause of inherited colorectal carcinoma (CRC). Testing all newly diagnosed CRC for MMR protein deficiency, known as universal testing, has recently emerged as the preferred approach to identify potential Lynch syndrome individuals. All newly diagnosed CRCs were screened for MMR protein expression by immunohistochemistry. A 2-step approach was used: PMS2 and MSH6 testing followed by the testing of the respective MMR protein partner if one of the proteins is lost. We retrospectively searched our pathology database for MMR protein expression results across a 5-year period (2012-2016) when universal testing was performed. Clinical and pathological data were extracted from the pathology report. A total of 2077 consecutive CRCs were tested for MMR protein expression. Mean age at diagnosis was 68.4 years. MMR protein deficiency was identified in 399 cases (19.2%). The vast majority of CRC with MLH1/PMS2 loss were diagnosed in patients older than 70 years (84%), most of them are likely to be secondary to sporadic *MLH1* methylation. MMR protein deficiency patterns suggestive of a defect in *MSH2, MSH6* or *PMS2* comprised 42 cases, of which 37 were found in individuals aged 50 years or older. CRCs with MSH2/MSH6 loss were most commonly found in patients older than 70 years (57%). In summary, universal testing for MMR protein deficiency in CRC identifies abnormal patterns of expression suggestive of Lynch syndrome in all age groups. Further studies are needed to demonstrate the actual rate of Lynch syndrome individuals identified from this initial screening.

## Introduction

In Australia, colorectal carcinoma (CRC) is the second most common cancer with a lifetime risk by age 75 years of 1 in 23 and is the second leading cause of death from cancer (1 in 48 by age 85).^1^ One way to reduce the mortality associated with CRC is the early detection of high risk individuals who then undergo additional screening. Lynch syndrome is the most common cause of inherited CRC, responsible for 3% of all incident cases.^2^ Lynch syndrome is an autosomal dominant disorder, defined by the identification of a pathogenic germline mutation in one of the DNA mismatch repair (MMR) genes *MLH1, MSH2, MSH6,* or *PMS2* or in *EPCAM,* a gene upstream of *MSH2.* Affected individuals have a cumulative risk of CRC at 70 years of 30-50% for MLH1 or MSH2 mutation carriers and of 10-20% for MSH6 or PMS2 mutation carriers.^3, 4^ Lynch syndrome individuals also have an increased risk of developing cancers of the endometrium, urinary tract, pancreas, hepatobiliary tract, stomach, small intestine and ovaries.

Among various screening strategies advocated, testing all newly diagnosed CRC for MMR protein expression by immunohistochemistry has emerged as the preferred approach to identify potential Lynch syndrome individuals. This reflex testing strategy, commonly named universal testing, is now recommended by multiple guidelines and professional medical organisations.^5–7^ The sensitivity of universal testing for detecting Lynch syndrome is close to 100%^8^ and, when compared with testing restricted to patients younger than 50 years at CRC diagnosis or fulfilling Bethesda criteria, can detect nearly twice as many individuals with Lynch syndrome with a favourable cost-effectiveness ratio per life-year saved.^9^

In this study, we investigated the proportion of CRC with abnormal MMR protein expression over a 5-year period when universal testing was performed in a community-based gastrointestinal pathology practice. Our aims were 1) to report the number of MMR deficient CRC cases using universal testing and 2) to identify possible Lynch syndrome individuals who would have been missed if a more restricted testing strategy was used.

## Materials and methods

Envoi Specialist Pathologists is a community-based histopathology practice in Brisbane, Queensland, Australia, that specialises in gastrointestinal pathology. Since 2012, all newly diagnosed CRCs have been screened for MMR protein expression by immunohistochemistry. Toretrospectively select the study cases, we searched our pathology database for MMR protein immunohistochemistry results using the term “MSH6” as key word, between January 1^st^ 2012 and December 31st 2016. Duplicate entries (those with immunohistochemistry performed on both biopsy and resection specimens) were removed. Clinical and pathological data were extracted from the pathology report, including age at diagnosis, sex, CRC location and TNM stage.

Identification of immunohistochemical expression of MMR proteins was performed in a 2-step approach. Firstly, each CRC was tested for PMS2 and MSH6 expression. CRCs with the normal retained expression of both PMS2 and MSH6 were classified as MMR-proficient. If loss of expression of either PMS2 or MSH6 was identified, the corresponding binding partner was subsequently tested (MLH1 for initial PMS2 loss, MSH2 for initial MSH6 loss). Immunohistochemistry was performed on Dako autostainer with ready to use antibodies from Dako (Carpinteria, CA, USA): clone ES05 for MLH1, clone FE11 for MSH2, clone EP51 for PMS2 and clone EP49 for MSH6. CRCs with loss of expression of at least one MMR protein in carcinoma cells were classified as MMR-deficient. In cases with synchronous or metachronous carcinomas, each tumour was separately tested.

## Results

A total of 2077 consecutive CRCs with MMR protein expression results were identified from 2016 patients. Mean age at CRC diagnosis was 68.4 (median 70; range 18-96 years). Females comprised 45.5% of incident cases. MMR protein deficiency was identified in 399 cases (19.2%). The detailed results of the abnormal immunohistochemical patterns are displayed in Table 1, stratified by age groups.

**Table 1.** Number of MMR-proficient and MMR-deficient colorectal carcinomas by pattern of MMR protein loss and age groups

|  | All ages | <50 years | 50-59 years | 60-69 years | ≥70 years |
| --- | --- | --- | --- | --- | --- |
| **MMR proficient** | 1678 | 171 (10%) | 274 (16%) | 489 (29%) | 744 (44%) |
| **MLH1/PMS2** | 357 | 13 (4%) | 11 (3%) | 34 (9%) | 299 (84%) |
| **MSH2/MSH6** | 23 | 2 (9%) | 1 (4%) | 7 (30%) | 13 (57%) |
| **MSH6** | 12 | 3 (25%) | 3 (25%) | 4 (33%) | 2 (17%) |
| **PMS2** | 7 | 2 (29%) | 0 (0%) | 2 (29%) | 3 (42%) |

**Figure 1.**
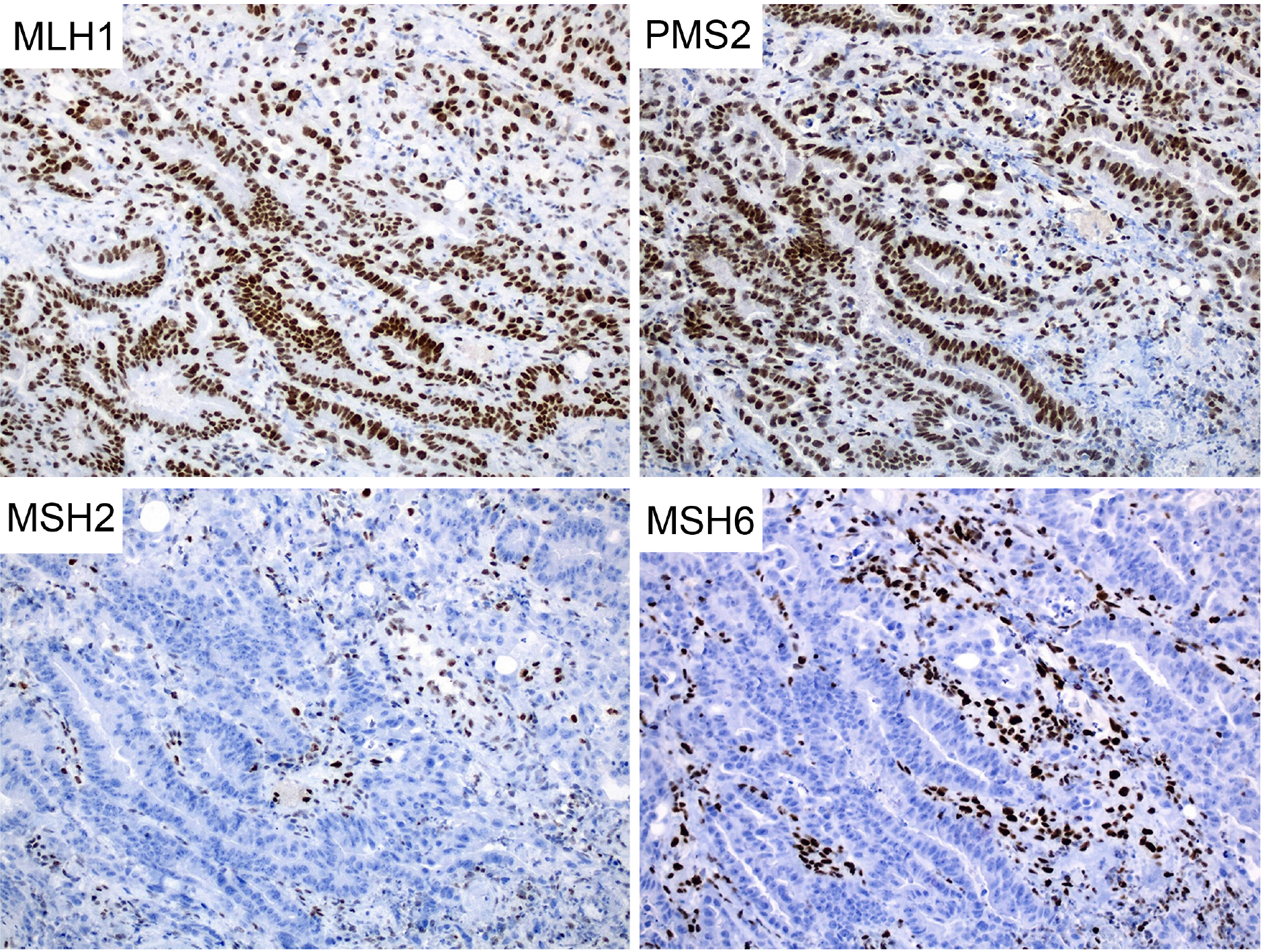
Colorectal carcinoma showing loss of immunohistochemical expression of MSH2 and MSH6, and retained expression of MLH1 and PMS2.

**Table 2.**
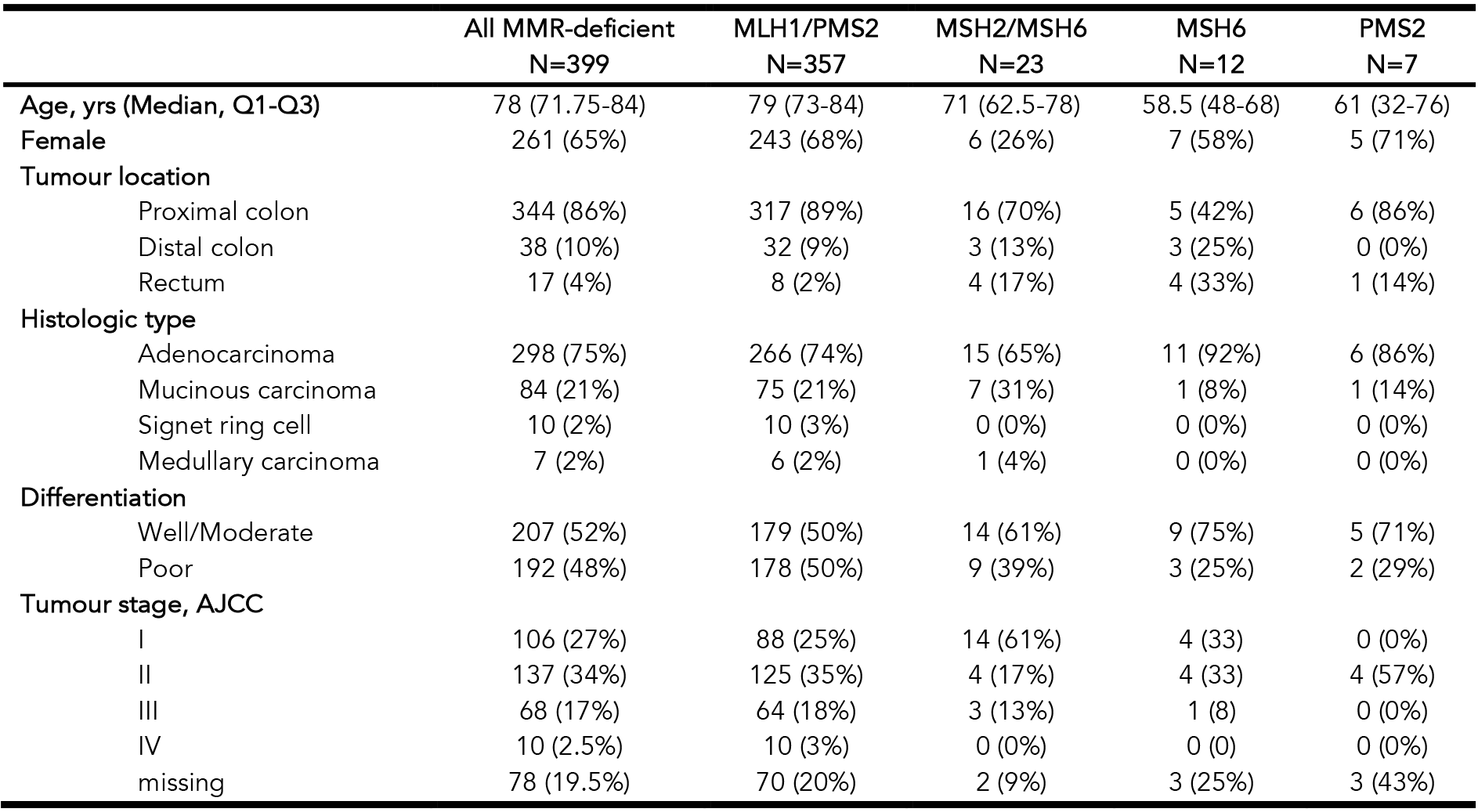
Clinicopathologic features of MMR-deficient colorectal carcinomas by pattern of MMR protein loss

The vast majority of CRC with MLH1/PMS2 loss were diagnosed in patients aged 70 years or older (84%). In 8 cases with MLH1/PMS2 loss, loss of MSH6 expression was also present. MMR protein deficiency patterns suggestive of a defect in MSH2, MSH6 or PMS2 were seen in 42 cases, of which 35 were found in individuals aged 50 years or older (Figure 1). CRCs with MSH2/MSH6 loss were most commonly found in patients older than 70 years (57%). The clinicopathological features of MMR-deficient cases are listed in Table 2. CRCs with MLH1/PMS2 loss were diagnosed more frequently in the proximal colon of females at a median age of 79 years. Cases with MSH2/MSH6 loss were characterized by male predominance, older age at diagnosis (median 71 years) compared with MSH6 loss or PMS2 loss, highest frequency of mucinous carcinoma (31%) and of stage I disease (61%). The rectum was most frequently involved in MSH6-deficient cases, compared with other MMR-deficient cases.

## Discussion

This study has investigated the incidence of MMR deficiency in a consecutive series of CRC that underwent universal testing by immunohistochemistry from a single Australian pathology practice in Brisbane, Queensland. We found that loss of one or more MMR protein by immunohistochemistry was seen in 19.2% of cases, with the vast majority being cases with loss of MLH1 and its binding partner PMS2 (89.5% of all MMR deficient cases). Over 80% of all cases of MMR deficiency were seen in patients over the age of 70 years, including 57% of cases with MSH2/MSH6 loss. As noted in other studies, the majority of MMR-deficient CRC occurred in the proximal colon and were low stage at diagnosis.^10^

This rate of MMR deficiency is higher than the 10-15% reported in most studies. One possible explanation could be a selection bias in our population. The average annual CRC incidence at Envoi Specialist Pathologists laboratory between 2012 and 2016 was 415 representing approximately 2.7% of the 15,151 Australians diagnosed with a CRC in 2011.^1^ From an estimated CRC incidence in Queensland around 3,000 in the same period of time, our population comprises 13.8% of all newly diagnosed CRC in this state, with a similar median age at diagnosis of 70 years as the one reported for the entire state of Queensland.^11^ The female proportion of 45.5% in our population was not different from the one in the entire Australian population (44.9%, P = 0.57). These findings support that our population is representative from the entire Australian population diagnosed with a CRC and argue against a selection from an older population with higher proportion of females who are more likely to develop MMR-deficient CRC with *MLH1* methylation. Two population-based studies have data on MMR deficiency in CRC. The Melbourne Collaborative Cohort Study is an Australian prospective cohort study which recruited Melbourne residents from 1990.^12^ In this cohort, the rate of MMR-deficiency was 15% of 740 incident CRC cases in a population with a mean age of 69 years at diagnosis and a female proportion of 46.2%. ^10^ This lower rate may at least be partially explained by a high proportion (25%) of individuals recruited in this study born in Mediterranean countries where the proportion of MMR deficiency in CRC has been reported to be <10%.^13, 14^ In another Australian study from the Royal North Shore Hospital in Sydney, 19.5% of 1426 CRCs diagnosed between 2004 and 2009 had MMR deficiency, a rate similar to our series. ^15^ Possible other causes for the high rate of MMR deficiency in our study include lifestyle factors such as smoking and dietary habits (red meat consumption, cooking practice including barbecuing) that have been reported to be associated with microsatellite instability.^16, 17^

Until recently, the recommendation for pathologists was to request MMR protein immunohistochemistry if patients meet the Amsterdam criteria or the revised Bethesda criteria. These clinical criteria include age at CRC diagnosis younger than 50 years and family history of Lynch syndrome-associated malignancies. In practice, pathologists are rarely aware of any delailed family history to apply these criteria when reporting a biopsy or a surgical specimen with CRC. Age at diagnosis was therefore the most common trigger to perform MMR protein immunohistochemistry. In our study, when compared with age restricted testing strategies, 35 of 42 cases (83%) with a MMR-deficient CRC suggestive of Lynch syndrome (tumours showing MSH2/MSH6 loss, MSH6 loss or PMS2 loss) would have been missed if testing was only performed for patients younger than 50 years at diagnosis. If the age restritction was 70 years, 18 patients (43%) would have been missed.

There was also a vast increase in the incidence of MLH1 deficiency in patients over 70 years, most of these are likely to be secondary to somatic *MLH1* promotor methylation. For CRC with loss of MLH1/PMS2 staining by immunohistochemistry, testing for the somatic *BRAF*^*V600E*^ mutation is recommended to help separate sporadic from Lynch syndrome-associated CRC. Mutation testing for *BRAF*^*V600E*^ is usually offered to patients under the age of 70 as nearly all MLH1-deficient CRC in patients over 70 are caused by *MLH1* methylation, not Lynch syndrome. If *BRAF*^*V600E*^ mutation is detected, the tumour is assumed to be MMR deficient due to *MLH1* methylation and no further investigations are performed unless there is a strong clinical suspicion of Lynch syndrome (young age or strong family history suggestive of Lynch syndrome). If *BRAF*^*V600E*^ mutation is not detected in the tumour, genetic counselling is recommended with germline testing for a mutation in *MLH1,* followed by *PMS2* if no mutation in *MLH1* is found. Prior to germline mutation testing, somatic methylation analysis of the *MLH1* gene can be performed to reduce the referral rate for genetic counselling if *MLH1* methylation is detected.^18^ Unfortunately, there is currently no Medicare Benefits Schedule (MBS) rebate for *BRAF*^*V600E*^ mutation testing, *MLH1* methylation status or for germline mutation testing in Australia, resulting in a high and increasing rate of unnecessary referral for genetic counselling now that universal testing has become more widely implemented.

Up to 70% of CRC demonstrating MMR-deficiency suggestive of Lynch syndrome do not have an MMR gene mutation identified by standard genetic testing approaches.^19^ These cases of so called Lynch-like syndrome or suspected Lynch syndrome can be caused by a false abnormal MMR immunostaining result, a hidden germline MMR gene mutation not detected by current sequencing methods, a mutation in other genes causing MMR deficiency, biallelic somatic MMR gene mutation or somatic cell mosaicism. From the two largest studies on Lynch-like cases, 50-70% of those were caused by biallelic somatic mutation that can be found in any of the 4 MMR genes.^20, 21^ Tumour testing for MMR gene mutation is likely to become the next step in the work up of Lynch-like cases to further exclude patients and their relatives from unnecessary Lynch syndrome surveillance.

Beyond screening for Lynch syndrome, MMR protein analysis provides important prognostic and predictive information for the management of CRC patients. MMR deficiency is an established good prognostic factor and predictor to poor response to 5-Fluorouracil therapy.^22^ *BRAF* mutation is associated with a poor outcome.^23^ The MMR status is also an essential predictor for the response to inhibitors of checkpoint proteins such as PD-1 and PD-L1. Patients with an advanced BRAF-mutated CRC will be have a different first-line therapy from those with *BRAF*-wildtype CRC and may now be enrolled in clinical trials using specific BRAF-inhibitors. For all these reasons, MMR protein status along with *KRAS, NRAS* and *BRAF* mutation status are now considered as essential components of CRC diagnosis in the new 8^th^ edition of the AJCC Colorectal cancer TNM staging system and are recommended to be tested for by the American Society for Clinical Pathology, College of American Pathologists, Association for Molecular Pathology, and American Society of Clinical Oncology.^24^

This study was limited by the absence of *BRAF* mutation testing or *MLH1* methylation analysis in all cases of MLH1 loss by immunohistochemistry. No germline mutation testing result was available for patients with a MMR deficiency pattern suggestive of Lynch syndrome.

In summary, universal testing for MMR protein deficiency in CRC identifies abnormal patterns of expression suggestive of Lynch syndrome in all age groups, including many in those excluded by current guidelines. Further studies are needed to demonstrate the actual rate of Lynch syndrome individuals identified from this initial screening.

